# SigBridgeR: An Integrative Framework and Toolkit for Comprehensive Screening and Benchmarking of Phenotype-Associated Cell Subpopulations in Single-Cell Transcriptomics

**DOI:** 10.64898/2026.05.08.723458

**Authors:** Yuxi Yang, Zeyu Yan, Hongqian Qian, Chunyang Wang, Liping Du, Yun Peng, Xiaoyun Bu, Jian-Guo Zhou, Shixiang Wang

**Affiliations:** Department of Biomedical Informatics, School of Life Sciences, Central South University, Changsha, China; Department of Urology, Peking University People’s hospital, Beijing, China; Department of Colorectal Surgery, Hunan Cancer Hospital/The Affiliated Cancer Hospital of Xiangya School of Medicine, Central South University; Department of Oncology, The Second Affiliated Hospital of Zunyi Medical University, Zunyi, China

**Keywords:** Single-cell RNA sequencing, phenotype-associated subpopulation, bulk transcriptomics, computational benchmarking, R package, precision medicine

## Abstract

Single-cell RNA sequencing has revolutionized our understanding of cellular heterogeneity, yet linking specific cell subpopulations to clinically relevant phenotypes remains a persistent challenge. Although multiple computational methods have been developed to bridge this gap, they are typically implemented as standalone packages with heterogeneous preprocessing pipelines, incompatible parameter conventions, and divergent output formats, thereby hindering rigorous cross-method benchmarking and reproducible multi-method workflows. Here, we present SigBridgeR, an extensible R framework and comprehensive toolkit that currently unifies eight state-of-the-art phenotype-associated cell screening algorithms within consistent workflows. We conducted a systematic benchmarking study across four cancer types HER2-positive breast cancer, triple-negative breast cancer, lung adenocarcinoma, and ovarian cancer using both binary phenotypes and patient survival endpoints. Our evaluation incorporated positive and negative control assessments based on differentially expressed genes and randomly selected marker panels, alongside quantitative accuracy comparisons using ground-truth cell labels. Building upon these insights, SigBridgeR provides standardized preprocessing for scRNA-seq and bulk transcriptomic data, unified algorithmic interfaces through a registry-based architecture, ensemble analysis via weighted voting, and comprehensive visualization utilities for multi-method comparison. By lowering technical barriers and promoting methodological standardization, SigBridgeR facilitates reliable discovery of phenotype-relevant cell subpopulations and enhances the translational potential of single-cell omics research.

## 1. Introduction

The advent of single-cell RNA sequencing (scRNA-seq) technologies has fundamentally reshaped our understanding of cellular heterogeneity within complex biological systems, offering unprecedented resolution to dissect tissue architecture, cellular states, and dynamic processes in both health and disease states^1–3^. However, despite these advances, translating high-resolution single-cell atlases into clinically actionable insights continues to present substantial methodological and analytical hurdles. A particularly salient limitation lies in the difficulty of establishing direct causal or associative links between specific cell subpopulations and macroscopic clinical phenotypes, such as patient survival outcomes, treatment responses, or disease progression trajectories^4,5^.

To address this gap, a growing family of computational methods—often termed phenotype-associated subpopulation identification or bulk-guided single-cell analysis approaches—has emerged over the past several years. These methods leverage the complementary strengths of scRNA-seq and bulk RNA sequencing (bulk RNA-seq) data: the former provides fine-grained cellular resolution, while the latter offers scalable, clinically annotated transcriptomic profiles from large patient cohorts. Representative tools include Scissor^6^, DEGAS^7^, scAB^8^, LP_SGL^9^, SCIPAC^10^, PIPET^11^, scPAS^12^ and scPP^13^. While each method introduces innovative algorithmic paradigms to link single-cell profiles with clinical phenotypes, they are typically implemented in separate software packages with inconsistent preprocessing pipelines, incompatible parameter conventions, and divergent output structures. These heterogeneities pose substantial barriers to rigorous comparative benchmarking, cross-method validation, ensemble modeling, and reproducible multi-method workflows in single-cell genomics research.

To overcome these limitations, we developed SigBridgeR, a comprehensive R toolkit that consolidates multiple state-of-the-art phenotype-associated cell screening algorithms within a unified, extensible framework. SigBridgeR incorporates standardized preprocessing modules for scRNA-seq data, bulk expression matrices, and diverse phenotype inputs—including binary traits, survival endpoints, and continuous variables—ensuring consistent data handling across all integrated methods. In this study, we describe the design and implementation of SigBridgeR, report a comprehensive benchmarking study evaluating eight algorithms across multiple cancer datasets and phenotype types, and demonstrate how the framework facilitates reproducible, translational single-cell omics research.

## 2. Results

### 2.1 Overview of Algorithms and Datasets for Evaluation

We systematically evaluated eight phenotype-associated cell screening algorithms spanning diverse methodological categories (**Table 1**): network-based regression (Scissor), deep transfer learning (DEGAS), non-negative matrix factorization (scAB), sparse group lasso with community detection (LP_SGL), quantitative association estimation (SCIPAC), phenotype-guided similarity matching (PIPET), network-regularized sparse regression with permutation testing (scPAS) and marker projection via AUCell enrichment (scPP) in select comparative analyses. These algorithms were applied to scRNA-seq datasets from four cancer indications—HER2-positive breast cancer, triple-negative breast cancer (TNBC), lung adenocarcinoma (LUAD), and ovarian cancer—paired with corresponding bulk RNA-seq datasets and phenotypic annotations (**Table S1**).

**Table 1:**
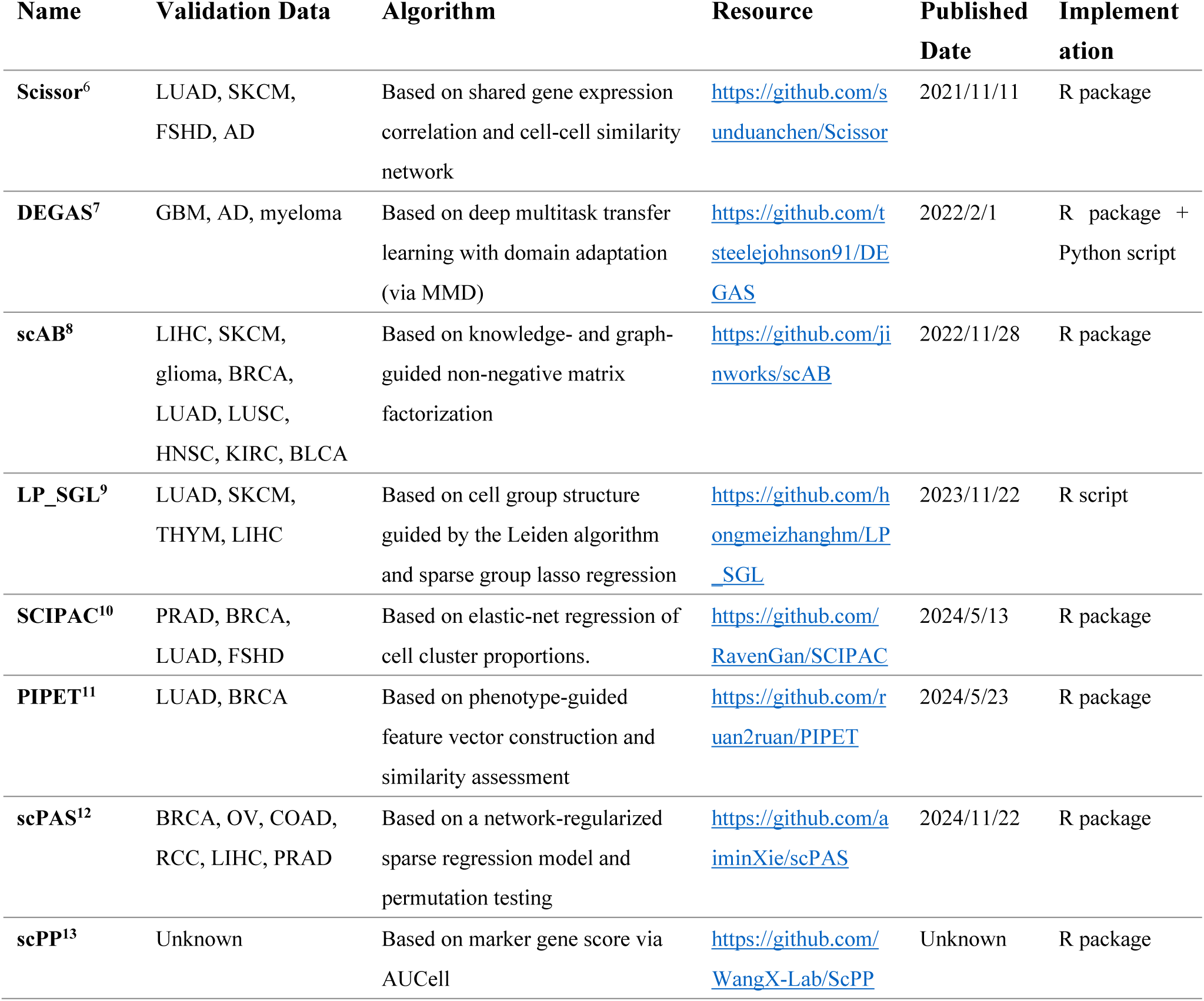
Phenotype-Associated Cell Screening Algorithms Evaluated in This Study.

The experimental design encompassed two primary phenotype categories: binary classifications (e.g., tumor vs. normal, recurrent vs. non-recurrent, high-grade vs. low-grade) and patient survival endpoints (overall survival time and status). Binary phenotypes were available for GSE42568 (tumor/normal), GSE162228 (relapse/non-relapse), TCGA-BRCA (tumor/normal), GSE9891 (malignant/low malignant potential), GSE140082 (high-grade/low-grade), and TCGA-LUAD (tumor/normal). Survival phenotypes were applied uniformly across all datasets where bulk RNA-seq and clinical follow-up data were co-available. The heterogeneity in phenotype definitions across datasets reflects real-world variation in clinical annotation.

All algorithms assigned labels to individual cells following execution. We harmonized output labels across methods into a unified nomenclature: Positive denoted cells positively associated with the phenotype (e.g., tumor-associated, poor survival); Negative denoted cells inversely associated; Neutral indicated weak or nonsignificant associations; and Other represented the collective set of non-Positive cells in two-class algorithms. Scissor, scPAS, scPP, SCIPAC, PIPET, and LP_SGL produced three-class outputs (Positive/Negative/Neutral), whereas scAB and DEGAS generated two-class outputs (Positive/Other). DEGAS required post hoc thresholding (score differential > 0.4 and top 5% ranking) to derive class labels, as its native output comprises continuous correlation scores rather than discrete assignments (see **<u>Methods</u>**).

### 2.2 Heterogeneity of Identifying Phenotype-Associated Cell Subpopulations

Across all dataset-algorithm-phenotype combinations, the identified Positive cell subpopulations exhibited marked heterogeneity in both quantity and distribution. UMAP visualization of screened subpopulations demonstrated that Positive cells frequently clustered within specific transcriptional neighborhoods, although substantial dispersion was observed for algorithms with less stringent selection criteria (**Figure 1A**). Under randomly selected markers from bulk RNA-seq data, the ssGSEA scores of different screening groups show no significant bias, demonstrating that the phenotype-associated cells identified by each algorithm are minimally affected by the data themselves or by random factors. Meanwhile, ssGSEA scoring based on bulk-derived positive markers confirmed that Positive subpopulations generally exhibited elevated enrichment scores relative to Negative, Neutral, or Other cells. However, the magnitude and specificity of enrichment varied considerably across algorithms, In TNBC, algorithmic behavior broadly paralleled results in HER2-positive cancers, though with notable differences in Positive cell fractions attributable to the distinct tumor microenvironment composition of triple-negative disease. LUAD and ovarian cancer analyses similarly revealed algorithm-specific sensitivities: LP_SGL frequently allocated substantial cell fractions to the Neutral category, consistent with its community-structure regularization that moderates extreme assignments, whereas scAB tended toward more binary-like partitions due to its matrix factorization objective.

**Figure 1:**
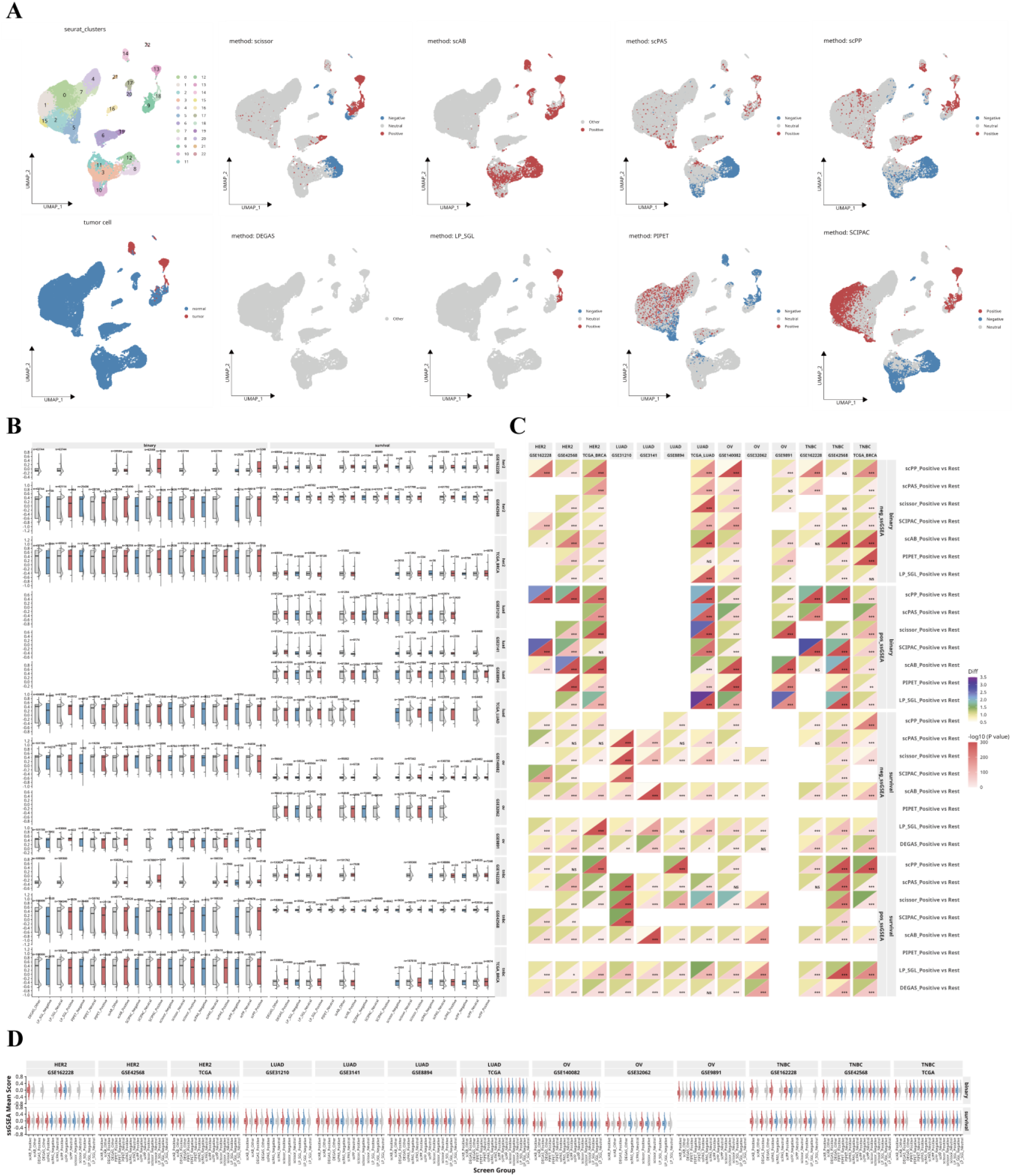
Overview of Phenotype-associated Cell Subpopulation Across Datasets. **(A)** Using the single-cell RNA-seq dataset, corresponding bulk RNA-seq dataset and phenotype, multiple algorithms were applied to identify phenotype-associated cell subpopulations. Exemplified by using LUAD dataset under binary phenotype. **(B)** Negative control experiments show the distribution of the average ssGSEA score per run across 100 repetitions for each screening group when using random markers from bulk RNA-seq expression data. **(C–D)** Positive control experiments using significant markers derived from bulk RNA-seq data show, under different data configurations: (C) the distribution of ssGSEA scores for each screening group, and (D) the significance of ssGSEA scores for the Positive subgroup. Diff is calculated from the average ssGSEA scores of the two groups.

### 2.3 Positive and Negative Control Assessments

Positive control experiments validated the biological relevance of identified subpopulations across most algorithm-dataset combinations (**Figure 1B, C**). Using the top 20 bulk-derived differentially expressed genes (DEG) as phenotype-associated markers, ssGSEA scores in the Positive group were significantly elevated relative to the most of Rest group of successful screenings. Scissor and LP_SGL demonstrated the most consistent positive control validation, with significant enrichment ratios and low p-values across breast cancer, LUAD, and ovarian cancer datasets under both binary and survival phenotypes. Notably, LP_SGL exhibited the best screening performance under binary phenotypes and also performed well under survival phenotypes, but encountered memory allocation failures on the largest cohorts (>50,000 cells), precluding assessment in 22.2% (2/9) of attempted screenings. scPAS and scPP exhibited strong positive control performance in most tested scenarios but occasionally produced nonsignificant ssGSEA separations in datasets with limited bulk-scRNA-seq gene overlap or weak phenotype signals. These algorithms have demonstrated good cross-data accuracy and robustness.

SCIPAC demonstrated excellent accuracy under the binary phenotype; however, under the survival phenotype, it failed to find Positive cell subpopulations in the majority of datasets (6/9, 66.7%). DEGAS positive control performance was generally strong when sufficient training epochs were allocated under survival phenotype; however, under binary phenotype screening, DEGAS failed to capture the corresponding positive cells in all data groups (**Figure 1B, C**). This may be due, on one hand, to the relatively large threshold we selected (see **<u>Methods</u>**), and on the other hand, to the possibility that DEGAS indeed performs poorly under binary phenotype screening. Under the binary phenotype, PIPET was generally able to identify Positive subpopulations that were significantly distinct from other cells; however, the ssGSEA scores of these cells were hardly distinguishable from those of other cells. Under the survival phenotype, PIPET generally failed to identify marker genes associated with the phenotype, resulting in no detection of phenotype-related Positive cell subpopulations. Overall, these algorithms only demonstrated good accuracy under certain circumstances.

Negative control experiments, employing randomly selected 20-gene marker panels across 100 replicates, provided critical validation of result specificity. Anderson-Darling tests confirmed that the distributions of mean ssGSEA scores for randomly selected markers showed no significant difference (AD statistic range: 0.0521–0.514, all p > 0.9), demonstrating that stochastic factors in ssGSEA did not significantly affect the conclusions of the positive control experiments. Furthermore, none of the algorithms exhibited methodological bias toward highly expressed genes.

### 2.4 Benchmarking Computational Performance

Computational performance profiling on the ovarian cancer dataset (**Figure 4**) revealed order-of-magnitude differences in runtime and memory requirements across algorithms. It should be noted that scPP and PIPET include marker gene selection in their core algorithm, so the number of marker genes obtained from the data significantly impacts computation time and memory usage. DEGAS imposed the heaviest computational burden, with training times scaling steeply with cell count regardless of dataset size due to its neural network architecture. LP_SGL’s reliance on the SGL package (v1.3)—which has not been actively maintained—resulted in memory errors on datasets exceeding approximately 50,000 cells, representing a critical limitation for contemporary large-scale single-cell atlases (**Figure 4B**).

**Figure 4:**
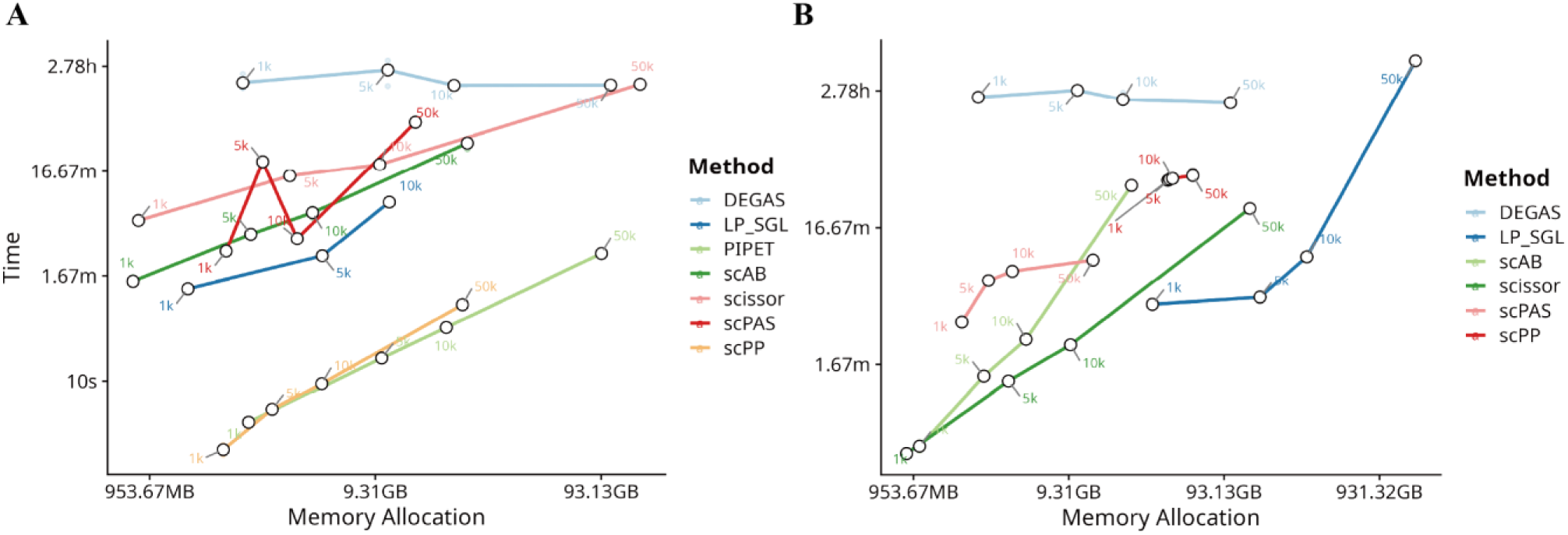
Computational performance and runtime of various single cell level phenotype-associated subpopulation screening algorithms under (A) binary phenotype data and (B) patient survival, respectively, evaluated on a single-cell ovarian cancer dataset GSE165897 paired with bulk RNA-seq dataset GSE140082. Data scale indicated on each curve represents the number of cells in the single-cell dataset.

### 2.5 SigBridgeR: A Unified Framework for Multi-Algorithm Screening

Motivated by the observed heterogeneity in algorithmic performance, parameter sensitivity, and computational requirements, we developed SigBridgeR as an integrative solution for standardized, reproducible phenotype-associated cell screening. The package architecture comprises three principal functional modules (**Figure 5**).

**Figure 5:**
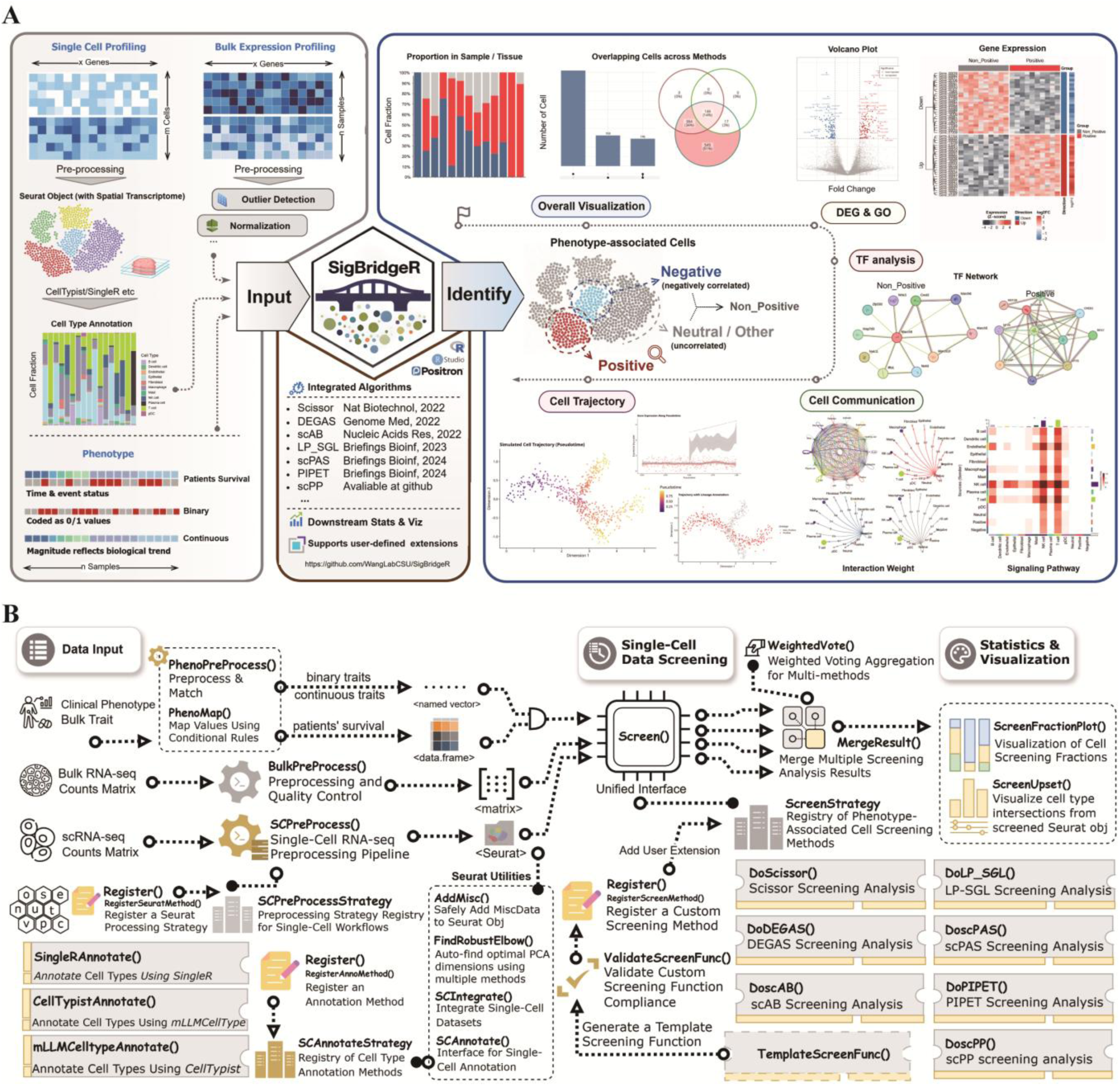
SigBridgeR: A Framework for Single-cell Level Phenotype-associated Cell Population Algorithms. **(A)** Workflow and functional roles of the SigBridgeR in biological analysis. **(B)** The function design and package structure of SigBridgeR

The preprocessing module provides standardized pipelines (**Figure 5**) for scRNA-seq data (SCPreProcess), bulk RNA-seq data (BulkPreProcess), and phenotype formatting (PhenoPreProcess), with automated handling of gene symbol conversions, missing value imputation, and cross-platform normalization. Special accommodations are included for preprocessed datasets (e.g., Scanpy-normalized data) through optional skipping of redundant normalization steps.

The screening module implements the registry-based algorithm invocation system (**Figure 5B**). Users call the unified Screen() function with dataset and algorithm specifications; the registry dispatches to the appropriate method while enforcing consistent input/output formats. Eight algorithms are currently registered (Scissor, scAB, scPAS, scPP, DEGAS, LP_SGL, PIPET, SCIPAC), with TiRank^14^ integration planned for the upcoming v3.7 release. The registry architecture permits straightforward extension through the Register() interface. A worked example of registering a custom algorithm is provided with detailed documentation (see **<u>Methods</u>**).

The integration and visualization module offers utilities for multi-method result comparison and ensemble analysis. MergeResult() consolidates outputs from multiple algorithms into unified data structures amenable to cross-method comparison. WeightedVote() implements ensemble classification by aggregating algorithmic labels through user-specified or performance-derived weighting schemes. ScreenFractionPlot() and ScreenUpset() generate visualizations of cell assignment overlaps and intersections across methods. Additionally, built-in cell annotation functions interface with SingleR, mLLMCelltype, and CellTypist to facilitate biological interpretation of identified subpopulations, and custom annotation methods are also supported.

SigBridgeR’s performance optimizations include data.table-based matrix operations, Rcpp-implemented bottleneck functions for scAB, SCIPAC, and parallel computation support via future/furrr. Benchmarking within SigBridgeR demonstrated 2- to 5-fold runtime reductions for optimized algorithms relative to their original implementations (**Figure 4**), with full numerical reproducibility verified across test suites.

## 3. Discussion

This study represents a comprehensive benchmark for phenotype-associated cell subpopulation screening algorithms, evaluating eight distinct methodological approaches across multiple cancer types, bulk RNA-seq pairings, and phenotype categories. Our findings reveal a landscape of considerable heterogeneity in algorithmic performance, robustness, and computational characteristics, underscoring both the wealth of available analytical options and the critical need for standardized integration frameworks such as SigBridgeR.

A primary observation from our benchmarking concerns the differential strengths and limitations of individual algorithms. Scissor and scAB emerged as the most consistently robust methods across datasets, maintaining reliable positive control validation and reasonable accuracy irrespective of bulk data source or phenotype type. Their shared reliance on regularized regression frameworks—network-based correlation optimization in Scissor and graph-guided matrix factorization in scAB—appears to confer resilience against the technical variability inherent in cross-platform transcriptomic integration. Furthermore, both methods support automated parameter iteration, substantially reducing user decision burden and promoting reproducible defaults. These characteristics position Scissor and scAB as dependable first-choice methods for exploratory analyses, particularly when ground-truth labels are unavailable for performance calibration.

LP_SGL and SCIPAC, despite achieving among the highest accuracy metrics when computable—consistent with the original report that they outperformed Scissor and scAB—showed less than satisfactory robustness across different data and computing platforms. In our comparative analysis, LP_SGL could barely handle datasets exceeding approximately 50,000 cells (**Figure 4B**). This limitation is particularly concerning given the increasing prevalence of large-scale single-cell atlases^15–19^. Regarding SCIPAC, although it achieved competitive accuracy under binary phenotypes, it lost this advantage under survival phenotypes, failing to identify phenotype-associated Positive cells in most datasets (**Figure 1B, C**). This implies that LP_SGL and SCIPAC are only suitable when pursuing extreme biological significance: the former requires a small data scale, while the latter is not applicable to patient survival data.

DEGAS occupies a complementary niche, offering exceptional accuracy when properly configured but imposing substantial computational and expertise requirements. Its deep transfer learning architecture effectively captures nonlinear gene expression relationships that linear models may overlook, and its maximum mean discrepancy-based domain adaptation mechanism mitigates batch effects between bulk and single-cell data modalities. However, the expansive hyperparameter space—including network depth, regularization coefficients, batch sizes, and dropout rates—necessitates either extensive grid searches or acceptance of potentially suboptimal performance. In a comparative context, contemporary deep learning approaches for single-cell analysis, such as foundation model-based methods, have demonstrated remarkable capability in gene expression modeling and represent a future trend—an advantage unique to deep learning in complex tissue scenarios and large-scale data^14,20^.

scPAS and scPP demonstrated solid middle-tier performance with distinct operational profiles. scPAS’s network-regularized sparse regression effectively leverages gene-gene relationships from scRNA-seq data, and its permutation testing framework provides statistically grounded significance estimates. Its performance was generally stable across datasets, though with occasional failures to identify significant Positive subpopulations when bulk phenotype signals were weak. scPP’s marker-based approach is conceptually straightforward and computationally efficient, making it attractive for rapid screening. Nevertheless, its reliance on bulk-derived marker lists rendered it sensitive to differential expression analysis outcomes; in scenarios where, bulk comparisons yielded limited significant markers—frequently observed in our ovarian cancer analyses—scPP’s discriminative capacity diminished markedly.

PIPET exhibited variable performance among evaluated algorithms. Its phenotype-guided feature vector construction is theoretically appealing, as it directly incorporates multivariate phenotypic information rather than reducing phenotypes to binary or survival endpoints. However, our results indicate that PIPET’s effectiveness is critically dependent on obtaining sufficient differentially expressed genes from bulk data to build informative feature vectors. In multiple dataset configurations, particularly those with modest bulk sample sizes or subtle phenotype differences, PIPET extracted fewer than 20 marker genes—insufficient for reliable subpopulation identification. This sensitivity to bulk data quality suggests that PIPET may be most appropriate for well-powered bulk cohorts with strong phenotype-associated transcriptional signals.

The discordance in Positive cell assignments across algorithms using identical scRNA-seq data but different bulk RNA-seq pairings highlights an important but often underappreciated source of variation in phenotype-associated screening. Bulk RNA-seq datasets differ in platform, sample size, tissue processing, and clinical annotation quality, all of which influence the phenotype signal transmitted to single-cell inference. Our observations suggest that screening outcomes may reflect bulk data characteristics as much as intrinsic single-cell biology, emphasizing the value of multi-bulk validation and ensemble approaches that aggregate evidence across multiple bulk cohorts.

Several methodological limitations of our study warrant explicit acknowledgment. First, our ground-truth accuracy assessments relied on intrinsic dataset labels—tumor cell annotations for breast cancer and CNV status for LUAD—that provide biologically meaningful but imperfect reference standards. True phenotype-associated cells may not perfectly align with categorical labels; for instance, non-malignant stromal cells exhibiting activated stress programs might genuinely associate with poor survival despite lacking tumor genotypes^21,22^. Second, using the packages bench (v1.1.4), microbenchmark (v1.5.0), peakRAM (v1.0.3) and callr (v3.7.6) sequentially within the R process to monitor memory scheduling, our computational performance benchmarking on large datasets (>100,000 cells) was invariably constrained by memory monitoring limitations, and we observed system crashes without exception. We mitigated this through extrapolation from subsampled benchmarks, but precise large-scale performance characterization remains incomplete. Notably, LP_SGL relies on the legacy, unmaintained SGL (v1.3) package, which readily causes memory leaks in practice, as already evidenced in our detection process (**Figure 4B**). Third, our evaluation focused on cancer datasets with relatively well-defined phenotype endpoints; performance characteristics may differ in non-malignant contexts such as neurodegenerative disease, autoimmune disorders, or developmental processes where phenotype-associated signals are more diffuse. Fourth, the heterogeneous binary phenotype definitions across datasets (tumor/normal, relapse/non-relapse, grade) reflect real-world clinical annotation variation but may confound direct cross-cancer comparisons; our Discussion interpretations accordingly emphasize within-cancer patterns over cross-cancer rankings.

The registry-based architecture of SigBridgeR was conceived to address precisely the integration challenges our benchmarking revealed. By decoupling algorithm-specific implementations from core data handling and visualization infrastructure, SigBridgeR enables methodologists to contribute new screening algorithms through standardized registration interfaces without requiring package-level modifications. This extensibility is particularly valuable in a rapidly evolving field where novel algorithms continue to emerge—we have recently incorporated SIDISH and TiRank into the development branch. We anticipate that community contributions will progressively expand SigBridgeR’s algorithmic repertoire, while the standardized benchmarking workflows established here provide a template for rigorous performance validation of future entrants.

Looking forward, several development priorities merit emphasis. The integration of spatial transcriptomics data represents an immediate opportunity, as spatial context provides critical microenvironmental information that dissociated scRNA-seq cannot capture. Accordingly, we have added support for integrating spatial transcriptomics data and have provided corresponding documentation to guide users. Additionally, the development of adaptive ensemble weighting schemes—where algorithm contributions are dynamically calibrated based on dataset characteristics rather than fixed user specifications—could further enhance reliability. Incorporation of multimodal single-cell data (chromatin accessibility via scATAC-seq, protein abundance via CITE-seq, spatial morphology) alongside transcriptomics would enable richer phenotype association models that capture epigenetic and post-transcriptional regulatory layers.

In conclusion, our comprehensive benchmarking demonstrates that no single algorithm universally dominates phenotype-associated cell screening; rather, optimal method selection depends on dataset scale, phenotype type, available computational resources, and intended downstream analyses. Scissor and scAB offer the most consistent robustness for general-purpose screening, LP_SGL excels when dataset scale permits, and SCIPAC does well under binary phenotype. SigBridgeR addresses this complexity by providing a unified, extensible, and performance-optimized platform that consolidates diverse algorithms beneath standardized data handling, execution, and visualization layers, supported by ensemble analysis capabilities and community-extensible algorithm registries. By promoting reproducible multi-method workflows and lowering technical barriers to rigorous benchmarking, SigBridgeR facilitates reliable discovery of phenotype-relevant cell subpopulations and contributes to the translational application of single-cell omics in precision medicine.

## 4. Methods

### 4.2 Data Collection and Preprocessing

We curated and incorporated scRNA-seq and bulk RNA-seq datasets from four distinct malignancies: breast cancer (HER2-positive and triple-negative subtypes), lung adenocarcinoma (LUAD), and ovarian cancer (OV). For LUAD, single-cell expression data were obtained from GEO accession GSE123902, as compiled by Kang et al^23^. Bulk RNA expression data and associated clinical phenotypes were retrieved from PRECOG (https://precog.stanford.edu/): GEO datasets GSE3141^24^, GSE8894^25^, and GSE31210^26^, alongside TCGA-LUAD^27,28^ obtained through TCGAbiolinks (v2.37.1)^29^. For ovarian cancer, scRNA-seq data GSE165897^30^ and metadata were downloaded directly from GEO, while bulk expression datasets GSE9891^31^, GSE32062^32^, and GSE140082^33^ were retrieved using GEOquery (v2.76.0)^34^. For breast cancer, scRNA-seq data encompassing both HER2-positive and triple-negative subtypes were sourced from GSE161529^35^, with bulk RNA-seq datasets GSE42568^36^ and GSE162228^37^ obtained via GEOquery; TCGA-BRCA data were processed identically to TCGA-LUAD. Clinical survival information for TCGA cohorts was integrated from UCSC Xena^38^ via the UCSCXenaShiny platform (v2.2.0)^39^.

Single-cell data preprocessing followed the standard Seurat (v5.3.0)^40^ workflow. Quality control filtering removed lowly expressed genes and cells with excessive mitochondrial gene proportions. Normalization was performed using NormalizeData with default parameters. Highly variable genes were identified via the “vst” method, followed by scaling (ScaleData), principal component analysis (RunPCA), nearest-neighbor graph construction (FindNeighbors, using the first 16 principal components), and clustering (FindClusters). Nonlinear dimensionality reduction was subsequently applied through RunTSNE and RunUMAP for visualization. Notably, GSE123902 had been preprocessed with Scanpy; therefore, we applied the inverse log transformation (expm1) prior to Seurat object creation with min.cells set to 400. Given that Kang et al^16^. had already performed sequencing depth correction, mitochondrial filtering, low-quality cell removal, and doublet exclusion^23^, we omitted redundant quality control steps for this dataset and proceeded directly to downstream analyses.

Bulk RNA-seq data preprocessing varied by source. GEO-derived bulk datasets were processed using accompanying feature data to convert Ensembl identifiers to gene symbols. TCGA bulk data employed IDConverter (v0.3.5)^41^ for identifier mapping. The few missing values in GSE140082 were imputed using sample-wise median values. Raw counts of bulk data were normalized using log2(count+ 1). To maximize gene overlap with scRNA-seq data, bulk expression matrices were not subjected to additional filtering at the preprocessing stage; all integrated algorithms automatically intersected genes between single-cell and bulk datasets during execution.

Phenotype data handling required careful harmonization. Binary phenotypes were defined as follows: for TCGA-LUAD and TCGA-BRCA, primary solid tumors (sample code 01) were coded as 1 and normal tissue (code 11) as 0, with other sample types excluded; for GSE9891, malignant samples were coded as 1 and low malignant potential samples as 0; for GSE140082, high-grade tumors were coded as 1 and low-grade as 0; for GSE42568, breast cancer tissue was coded as 1 and normal tissue as 0; for GSE162228, recurrent tumors were coded as 1 and non-recurrent as 0. Survival phenotypes uniformly utilized overall survival time and status, with death coded as 1 and survival as 0 (**Table S2**). Bulk datasets lacking appropriate binary phenotypes (GSE3141, GSE8894, GSE31210 for LUAD; GSE32062 for OV) were excluded from binary phenotype analyses. The heterogeneous nature of these phenotype definitions across datasets reflects real-world variation in clinical annotation practices; we explicitly acknowledge this as a potential confounder in cross-cancer comparisons and address its implications in the **<u>Discussion</u>**.

### 4.2 Identification of Phenotype-Associated Cell Subpopulations

We evaluated eight open-source phenotype-associated cell screening algorithms spanning diverse methodological categories (**Table 1**). Scissor (v2.0)^6^ employs a network-based regression approach that quantifies similarity between single cells and bulk samples, subsequently optimizing a regression model on the correlation matrix against sample phenotypes to identify relevant subpopulations. scAB (v1.0)^8^ implements knowledge- and graph-guided non-negative matrix factorization (NMF) to detect multiresolution cell states with clinical significance. scPAS (v0.2)^12^ constructs a network-regularized sparse regression model integrating bulk expression profiles and gene-gene similarity networks from scRNA-seq data, followed by permutation testing for significance estimation. scPP (v0.0.0.9)^13^ identifies phenotype-associated marker genes from bulk data and evaluates their enrichment in single cells using AUCell scores, defining phenotype-positive cells through rank-based intersection. DEGAS (v1.0)^7^ utilizes deep multitask transfer learning with maximum mean discrepancy-based domain adaptation to transfer disease impressions from bulk-level patients to individual cells. LP_SGL^9^ incorporates cell group structure obtained via the Leiden algorithm into a sparse group lasso regression framework. PIPET (v0.1)^11^ generates feature vectors for each phenotype from bulk differentially expressed genes and identifies relevant cellular subpopulations through similarity assessment. SCIPAC (v0.1)^10^ clusters single cells, fits elastic net regression on bulk data to derive coefficients, computes quantitative cell-phenotype association strengths via a closed-form vector product, and assesses significance through bootstrap p-values.

Algorithm parameters were configured to balance default recommendations with analytical rigor. For DEGAS, cells were labeled as Positive if score differentials exceeded 0.4 and ranked in the top 5% under survival as phenotype, or if score differentials exceeded 0.7 under binary phenotype. Since patient stratification is not required^6–8,12^, binary labels offer better classification accuracy than survival data, so we adopted a larger threshold to obtain the nearly same proportion of Positive cells. All remaining cells were designated as Other. All other algorithm parameters were kept at developer-recommended defaults unless otherwise specified. PIPET only supports bulk-level phenotypic data labels but does not natively support patient stratification based on survival data. Therefore, we input patients’ survival status as binary labels.

### 4.3 Control Assessments and Validation Framework

To rigorously assess algorithmic performance, we established complementary positive and negative control frameworks. For positive controls, we extracted the top 20 differentially expressed genes (DEGs) from each bulk dataset, ranked by absolute fold change and p-value (prioritizing statistical significance), and used these as phenotype-associated markers for single-sample gene set enrichment analysis (ssGSEA). Mean ssGSEA scores were computed for each screened subpopulation. For three-class algorithms (Scissor, scPAS, scPP, PIPET, LP_SGL, SCIPAC), the enrichment ratio (designated diff) metric was calculated as the ratio of mean ssGSEA scores between the Positive group and the merged Negative/Neutral (designated Rest) group. For two-class algorithms (scAB, DEGAS), diff was computed as the ratio between Positive and Rest (Other) groups. Statistical significance was assessed via Wilcoxon rank-sum tests, with p < 0.05 denoted by *, p < 0.01 by **, and p < 0.001 by *; all other cases were marked as NS (non-significant).

For negative controls, we randomly selected 20 genes from the intersection of single-cell and bulk RNA-seq gene sets, repeated this sampling 1,00 times, and computed mean ssGSEA scores per screened subgroup for each replicate. Anderson-Darling k-sample tests (kSamples v1.2.12) were applied to validate the distributional differences across replicates. This procedure confirmed that observed ssGSEA patterns neither arose from stochastic gene selection nor introduced bias across various screened subgroups.

### 4.4 Accuracy and Precision Benchmarking

To evaluate algorithmic accuracy under ground-truth settings, we optimized key parameters through random sampling within specified ranges and assessed classification performance using intrinsic dataset labels. For TNBC and HER2-positive breast cancer, tumor cell annotations from GSE161529 served as ground truth, with Positive predictive value defined as the proportion of predicted Positive cells that were true tumor cells. For LUAD, copy-number variation (CNV) status (tumor vs. normal) derived from inferCNV served as the reference standard. Classification performance was quantified through F1-score, precision, and accuracy metrics. Parameter sweeps covered clinically relevant ranges: Scissor alpha from 0.001 to 0.95 with cutoff 0.05-0.5; scAB alpha/alpha_2 from {0.0005, 0.001, 0.005, 0.01, 0.05} with tred 0.5-5.0 and repeat_times 5-20; scPAS nfeature 500-5000 with imputation methods (None, KNN, ALRA); scPP probs 0-0.5 with Log2FC_cutoff 0-1; DEGAS architecture (DenseNet, Standard) with ff_depth 2-10, bag_depth 3-10, lamb1/lamb2/lamb3 2-10, scbatch_sz 50-500, patbatch_sz 25-100, hidden_feats 25-100 and do_prc 0.1-0.9; LP_SGL resolution 0-1 with nfold 5-20; PIPET distance metrics (cosine, pearson, spearman, kendall, euclidean, maximum) with nPerm 500-2500 and Log2FC 1.2-3; SCIPAC n_pc metrics 10-100, resolution 0.1-4, ela_net_alpha 0.1-1, hvg 500-5000, bt_size 10-100 with nfold 3-30. Default parameter configurations were included as reference benchmarks in all randomized trials.

### 4.5 Performance and Scalability Evaluation

Computational performance was evaluated using the ovarian cancer dataset (GSE165897 scRNA-seq paired with GSE140082 bulk RNA-seq) under both binary and survival phenotypes. Memory consumption and runtime were monitored using the bench package (v1.1.4), with gradient curves generated to depict scaling behavior across varying data sizes (1,000, 5,000, 10,000, and 50,000 cells). For datasets exceeding 100,000 cells, memory profiling encountered system limitations; performance metrics for large-scale data were therefore extrapolated from observed scaling trends. All benchmarking was conducted on a server with 937 GB RAM 96 physical CPU cores (192 logical threads), and a base frequency of 2.6 GHz to standardize hardware effects.

### 4.6 SigBridgeR Package Development

SigBridgeR was developed in R (v4.4.1) as an integrated toolkit providing unified scRNA-seq and bulk RNA-seq preprocessing, cell annotation, algorithm invocation, and visualization. The package employs Seurat (v5.3.0) for single-cell data management and exposes algorithmic interfaces through the central Screen() function. Visualization utilities leverage ggVennDiagram (v1.5.4)^42^ for overlap depiction, ggupset (v0.4.1)^43^ for intersection visualization, and patchwork (v1.3.2) for composite figure assembly. High-performance computing is facilitated through data.table (v1.18.2.1), matrixStats (v1.5.0), and cheapr (v1.5.0), with select algorithmic components rewritten in Rcpp (v1.1.1)^44^ and RcppArmadillo (v15.2.3.1)^45^ to enhance computational efficiency while preserving result reproducibility. The package adopts a registry-based extension architecture, enabling users to register custom preprocessing workflows, screening algorithms, and annotation methods through standardized interfaces (https://wanglabcsu.github.io/SigBridgeR/articles/Extending.html#extend-screening-methods). Random seeds were set (seed = 123) for all stochastic operations to ensure exact reproducibility. SigBridgeR is freely available at https://github.com/WangLabCSU/SigBridgeR and distributed under the GPL-3.0 license. A permanent software deposit with version-pinned dependencies is available via the release page.

## Data Availability

All single-cell and bulk RNA-seq datasets analyzed in this study are publicly available through Gene Expression Omnibus (GEO) and The Cancer Genome Atlas (TCGA) as detailed in Methods. benchmarking results, parameter optimization tables, and analysis scripts are deposited at https://github.com/WangLabCSU/paper_SigBridgeR. The SigBridgeR package source code, documentation, and vignettes are available at https://github.com/WangLabCSU/SigBridgeR.

## Contributors

YXY was responsible for software development and methodology. ZYY and LPD aided the development of PIPET extension. YXY, GJZ, and SW contributed to the conceptualization and study design. YXY, HQQ, CYW and SW performed data curation, formal analysis, and visualization. SW and GJZ provided project administration and resources. YXY and SW wrote the original draft. GJZ, SW supervised the study. All authors performed writing – review & editing.

## Acknowledgements

We are grateful for resources from the High-Performance Computing Center of Central South University, as well as to the Bioinformatics Platform of Furong Laboratory and the Bioinformatics Center of Xiangya Hospital, Central South University, for providing computational resources.

This study also used data generated by the TCGA Research Network (https://www.cancer.gov/tcga), which we thank for making these data publicly available.

